# Enteric nervous system regeneration and functional cure of experimental digestive Chagas disease with trypanocidal chemotherapy

**DOI:** 10.1101/2022.12.14.520395

**Authors:** Archie A. Khan, Harry C. Langston, Louis Walsh, Rebecca Roscoe, Shiromani Jayawardhana, Amanda F. Francisco, Martin C. Taylor, Conor J. McCann, John M. Kelly, Michael D. Lewis

## Abstract

Digestive Chagas disease (DCD) is an enteric neuropathy caused by *Trypanosoma cruzi* infection. There is a lack of evidence on the mechanism of pathogenesis and rationales for treatment. We used a mouse model that recapitulates key clinical manifestations to study how infection dynamics shape DCD pathology, and the impact of treatment with the front-line drug benznidazole. Curative treatment at 6 weeks post-infection resulted in sustained recovery of GI transit function, whereas sub-curative treatment led to infection relapse and gradual return of DCD symptoms. Neuro-immune gene expression profiles shifted from chronic inflammation to a tissue repair signature after cure, accompanied by increased glial cell activity and regenerative neurogenesis in the myenteric neuronal plexus. Delaying treatment until 24 weeks post-infection led to a partial reversal of the DCD phenotype, suggesting the accumulation of permanent tissue damage over the course of chronic infection. Our study shows that murine DCD pathogenesis is sustained by chronic *T. cruzi* infection and is not an inevitable consequence of acute stage denervation. The risk that irreversible enteric neuromuscular tissue damage and dysfunction will develop highlights the importance of prompt diagnosis and treatment. Finally, these findings support the concept of treating asymptomatic *T. cruzi* infected individuals with benznidazole to prevent DCD development.

## Introduction

Chagas disease (CD), or American trypanosomiasis, is a neglected tropical disease with a prevalence of 6.5 million cases, a burden of 10,000 deaths per year, 275, 000 DALYs and economic costs reaching US$7 billion per year (1, 2). The large majority of cases occur in endemic regions of Latin America, but there is a clear long-term trend of globalisation (3-5). CD is caused by *Trypanosoma cruzi*, a protozoan parasite, which is primarily transmitted to humans by blood-feeding insect vectors (triatomine bugs), but it can also be acquired congenitally or from contaminated blood transfusions, organ transplants and foodstuffs (6). Anti-parasitic treatment is limited to the nitroheterocyclic drugs, nifurtimox and benznidazole. Both have long dosing schedules and can cause significant toxicity (7, 8). Daily benznidazole treatment for 60 days is the current standard of care because side effects are considered less severe than for nifurtimox. Recent trial data show that reducing the duration of treatment to 2 weeks may be justified (9).

Upon transmission, *T. cruzi* invades target cells of diverse types and begins an approximately weekly cycle of replication, host cell lysis and dissemination. In most cases, adaptive immunity suppresses parasite numbers to very low levels; sterile clearance is considered rare (10, 11). Clinical manifestations affecting the heart and/or GI tract develop in around one third of chronically infected people. Benznidazole treatment is recommended for all acute, congenital and immunosuppression-related reactivation cases, as well as chronic infections in children and women of childbearing age (12). However, the evidence for the efficacy of benznidazole in terms of chronic disease progression and outcomes is limited. Treatment showed no significant benefit compared to placebo in terms of preventing death or disease progression in patients who already had symptomatic cardiac Chagas disease (8). There are no clinical or pre-clinical data on the impact of treatment on digestive Chagas disease (DCD) outcomes.

DCD is an enteric neuropathy characterised by progressive dilatation and dysfunction of sections of the GI tract (13, 14). Symptoms include achalasia, abdominal pain, constipation and faecaloma. Eventually, massive organ dilatation results in megasyndromes, usually of the colon and/or oesophagus. Dilatation is associated with loss of enteric neurons leading to peristaltic paralysis and smooth muscle hypertrophy. Options for DCD management are limited to palliative and surgical interventions (15), often implemented in emergency scenarios late in the disease course, with significant mortality risk (16).

DCD is thought to stem from collateral damage to enteric neurons caused by anti-parasitic inflammatory immune responses in the muscle wall of the affected region of the GI tract (17). Beyond this, the mechanism and kinetics of denervation, and therefore a rationale for treatment, are poorly defined. The inability to detect gut-resident parasites in chronic infections supported a model of acute phase damage unmasked by further ageing-related denervation (18). Molecular detection of *T. cruzi* DNA and inflammatory infiltrates in post-mortem and biopsy studies of human DCD circumstantially suggests that chronic parasite persistence may contribute to disease development (19-27). These data from late and terminal disease states are difficult to interpret in respect of relationships between pathogenesis and infection load or distribution over time. Experimental bioluminescence imaging and tissue PCR studies in mice showed that the GI tract is a major long-term reservoir of infection with diverse *T. cruzi* strains (28-32). This led to the development of a robust mouse model of DCD, which features significantly delayed GI transit associated with co-localised parasite persistence and enteric neuronal lesions in the wall of the large intestine (33). Here, we utilised this model to formally test the hypothesis that benznidazole-mediated cure of *T. cruzi* infection can either prevent DCD, or reduce its severity.

## Results

### Benznidazole-mediated cure of *T. cruzi* infection in the experimental DCD model

We have developed C3H/HeN mice infected with bioluminescent TcI-JR parasites as a model of chronic DCD (33)(Fig. 1). Subsets of mice were treated with benznidazole or vehicle at 6 weeks post-infection (wpi) (Fig. 1a, b). At this time, parasite loads are already in sharp decline as a result of adaptive immunity. *In vivo* bioluminescence imaging (BLI) showed that untreated mice transitioned to a stable, low-level chronic infection. In contrast, parasite loads in benznidazole treated mice became undetectable (Fig. 1b, 1c). This was corroborated by splenomegaly, low body weight and loss of GI mesenteric tissue at 36 wpi in the untreated infected group. In all cases, the read-outs reversed closer to control baseline after curative benznidazole treatment (Fig. 1d, Extended Data Fig. 1). *In vivo* and *ex vivo* BLI identified a subset of infected, BZ-treated mice (n = 11/27, 41%) in which the infection had relapsed (Fig. 1b, 1c, 1e). Of these, 5 (19 %) infections were only detectable by post-mortem *ex vivo* imaging of internal organs (Fig. 1e). Retrospective comparison of body weights and parasite loads showed there was no difference at the start of treatment between animals that were cured and those that relapsed (Extended Data Fig. 2). Relapse infections were most often localised to GI tissues, in contrast to the broadly disseminated pattern seen in the infected group (Fig. 1f, 1g). Of note, in the context of cardiac Chagas disease, relapse infections rarely localised to the heart (2/11, 18%), which was a site of frequent and high intensity parasitism in the untreated group (16/18, 89 %) (Fig. 1f, 1g). Overall, our findings show that benznidazole treatment at 6 wpi achieves 60 % parasitological cure in this experimental DCD model with infection relapse cases often localised to the GI tract.

**Figure 1:**
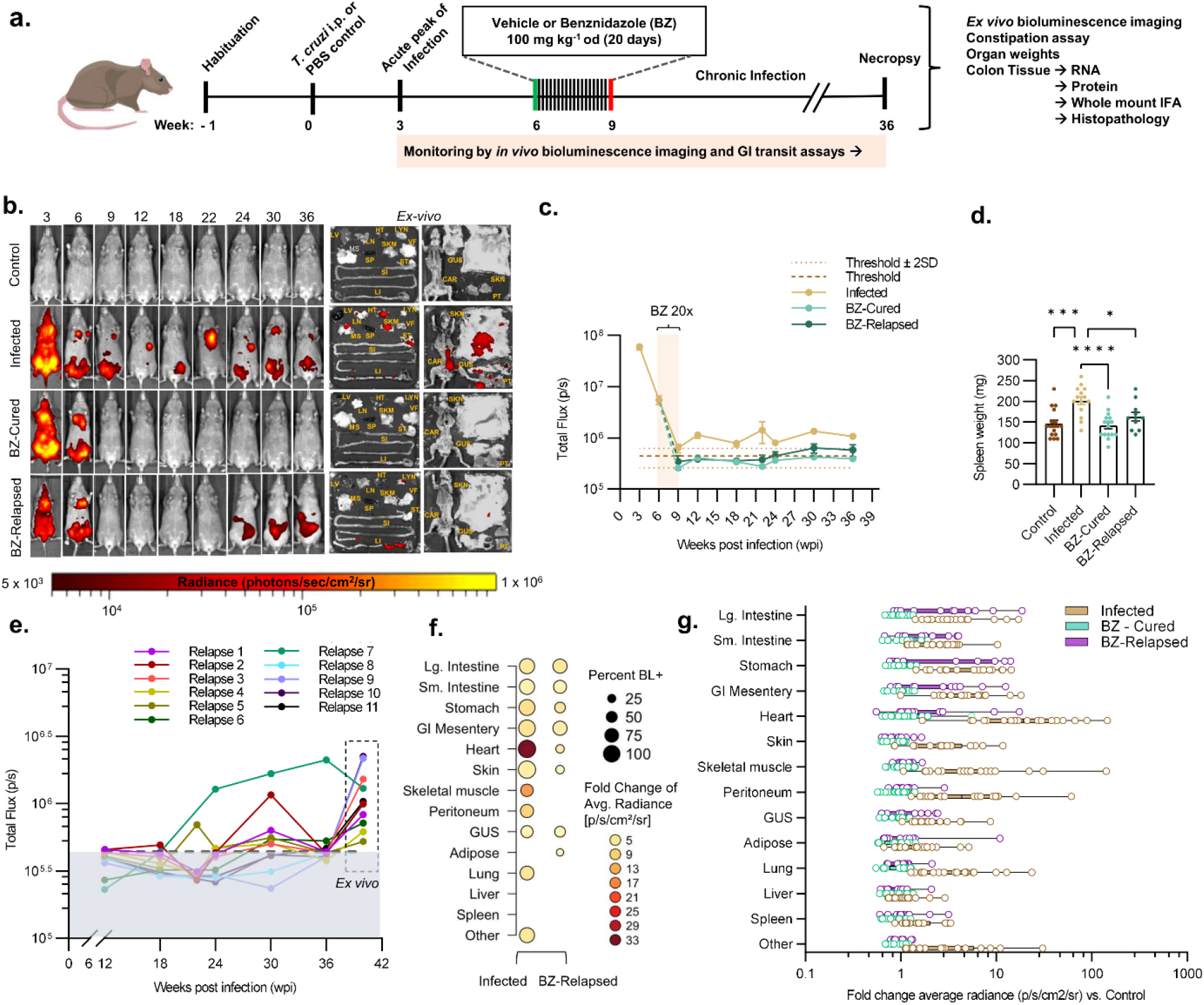
Evaluation of benznidazole treatment outcomes in a mouse model of digestive Chagas disease. **a**, Schematic representation of the experimental timeline. **b**, Representative *in vivo* bioluminescence images for each group of mice: uninfected controls (Control), *T. cruzi* infected and untreated (Infected), treated with benznidazole and parasitologically cured (BZ-Cured), and relapse infections after benznidazole treatment failure (BZ-Relapsed). *Ex vivo* bioluminescence images show parasite localisation in the indicated organs and tissue samples (liver-LV, lymph nodes -LYN, lungs - LN, gut mesenteric tissue - MS, heart - HT, spleen - SP, skeletal muscle - SKM, visceral fat - VF, stomach - ST, small intestine - SI, large intestine - LI, skin - SK, carcass - CAR, genitourinary system - GUS, and peritoneum - PT). Bioluminescence intensity is expressed on log-scale pseudocolour heat maps (range expressed in units of radiance). **c**, Line plots show course of infection as mean total bioluminescent flux (p/s) per mouse. Infected, *n* = 18, except *n* =10 at 9 and 22 wpi, *n* = 47 at 6 wpi and *n* = 50 at 3 wpi; BZ-cured, *n* = 16, except *n* = 17 at 12 and 18 wpi, *n* = 8 at 9 wpi and *n* = 7 at 22 wpi, and BZ- relapsed, *n* = 11, except *n* = 5 at 9 and 22 wpi. Dashed lines show thresholds: uninfected control mean auto-luminescence (dark brown dashed line) and mean ± 2SD (light brown dotted lines). **d**, Spleen weights at post-mortem of control (*n* = 17), infected (*n* = 14), BZ-Cured (*n* = 16) and BZ-Relapsed (*n* = 11) groups. **e**, Parasite bioluminescence intensity detected in individual BZ-Relapse mice after the end of benznidazole treatment (12 wpi). Data up to 36 wpi are from whole animal *in vivo* imaging and for 42 wpi are the sum of bioluminescent flux signals detected in organs and tissue samples imaged *ex vivo*. Dashed line shows uninfected control mean auto-luminescence threshold. **f**, Dot plot and **g**, box plot show relative *ex vivo* infection intensities in the indicated organs/tissue at 42 wpi. Data are expressed as the mean fold change in bioluminescent radiance vs. the uninfected control mean for groups (f) and individual mice (g). In dot plot **f**, the circle sizes indicate the percentage of mice with bioluminescence positive (BL^+^) signal and the circle colours show the infection intensity (fold change of mean bioluminescent radiance vs uninfected control). Infected, *n* = 18; except skin, GUS and other *n* = 17; BZ-Cured, *n* = 16 and BZ-Relapsed, *n* = 11 mice. Statistical significance was tested using one-way ANOVA followed by Tukey’s HSD test (\**P* < 0.05, \*\*\**P* < 0.001, **** *P* < 0.0001).

### Benznidazole treatment restores normal GI transit function associated with re-innervation

In the DCD model there is a highly significant delay in GI transit time in infected mice compared to uninfected controls ((33), Fig. 2a, 2b, 2c). Benznidazole chemotherapy rapidly reversed the transit delay phenotype to uninfected control baseline (Fig. 2b, 2c). Immediately prior to initiation of treatment, the mean GI transit time was 184 minutes in infected mice compared to 103 minutes in uninfected controls (Fig. 2b). In the untreated infection group, the delay remained significant, although it initially eased in line with immune-mediated parasite load reduction and then gradually worsened as the chronic phase progressed (Fig. 2b, 2c). Curative benznidazole treatment led to permanent restoration of normal transit times. Importantly, relapse infections were associated with the return of a significant transit delay, but this remained less severe than for the infected group (Fig. 2b, 2c, Extended Data Fig. 2). At the experiment end point, we analysed faecal retention in the colon after a period of fasting. This showed a clear constipation phenotype associated with an increased number of faecal pellets and weight in untreated infections. This was alleviated in benznidazole cured mice, but not in relapsed infections (Fig. 2d, 2e, 2f and Extended Data Fig. 1e). We also observed significant normalisation of caecum weight in cured mice (Extended Data Fig. 1c).

**Figure 2:**
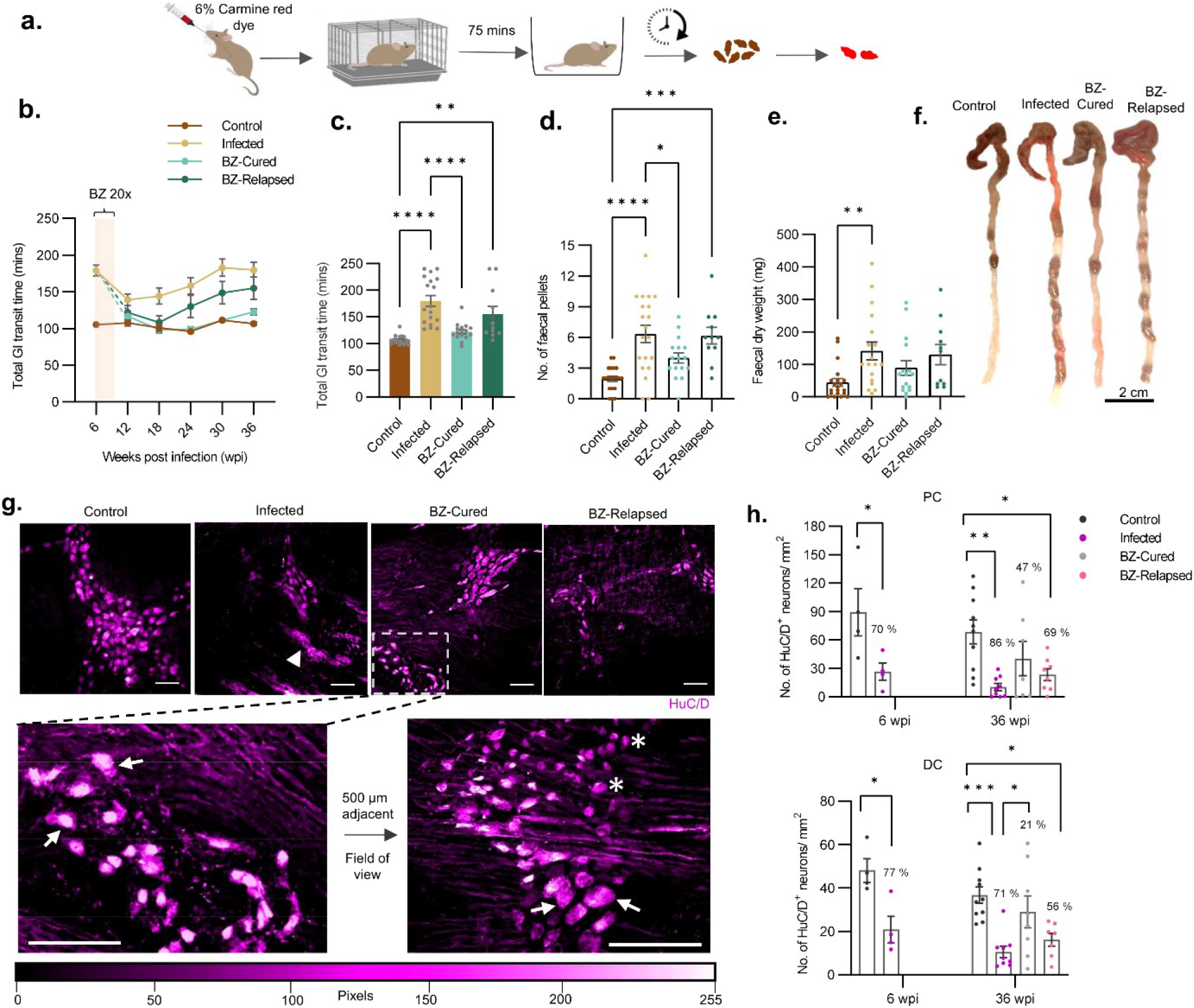
Benznidazole-mediated cure of infection reverses digestive Chagas disease phenotype and promotes ENS recovery. **a**, Schematic representation of the carmine red dye tracer method to measure transit time delay in mice. **b**, Line plots show change in mean total GI transit time over the course of 36 weeks of *T. cruzi* infection for infected (*n* = 18, except *n* = 46 at 6 weeks post-infection (wpi)), benznidazole (BZ) treated and cured (BZ-Cured; *n* = 16, except *n* = 17 at 12 and 18 wpi) and benznidazole treated and relapsed (BZ-Relapsed; *n* = 11) mice compared with uninfected controls (*n* = 20). Cream overlaid bar shows BZ treatment window (6-9 wpi, 20 daily doses). **c**, Bar plots show final GI transit times at 36 wpi for individual mice. Control *n* = 19, infected *n* = 18, BZ-Cured *n* = 16 and BZ-Relapsed *n* = 11. **d, e**, Bar plots show number of faecal pellets number (d) and dry weight (e) retained in the colon after 4 hours of fasting at 42 wpi for control (*n* = 20), infected (*n* = 18), BZ-Cured (*n* = 16) and BZ-Relapsed (*n* = 11) mice. **f**, Representative images of large intestines corresponding to the data in d and e. Scale bar = 2 cm. **g**, Representative immunofluorescence images of enteric neuronal cell bodies (HuC/D^+^, magenta) in the colon myenteric plexus from control, infected, BZ-cured and BZ-relapsed mice. White arrowhead indicates denervated ganglia (absence of defined soma). Magnified region of the BZ-Cured image shows intact control-like neuronal cell bodies, indicated by white arrows. In a 500 μm laterally adjacent region of the BZ-Cured ENS, a morphologically heterogeneous neuronal cell population is observed: white arrows indicate intact control-like neuron soma morphology and white stars indicate atypical neuron soma morphology. The magenta to white heat map scale corresponds to pixel fluorescence intensity. **h**, Bar plots show the densities of HuC/D-positive neurons in proximal and distal colon (PC, DC) before (6 wpi) and after BZ treatment (36 wpi) of control (*n* = 4 at 6 wpi, *n* = 10 at 36 wpi), infected (*n* = 4 at 6 wpi, *n* = 9 at 36 wpi), BZ-Cured (*n* = 7) and BZ-Relapsed (*n* = 8) groups. Data are expressed as mean ± SEM. Statistical significance was tested using one-way ANOVA followed by Tukey’s HSD test (\**P* < 0.05, \*\**P* < 0.01, \*\*\**P* < 0.001 **** *P* < 0.0001).

Given the central role of denervation in human DCD, we next evaluated the impact of infection and benznidazole-mediated treatment on the enteric nervous system (ENS). The number of neuronal cell bodies in the proximal colon myenteric plexus of infected-untreated mice was reduced by 70 % and 86% at 6 and 36 wpi respectively (Fig. 2g, 2h). The distal colon showed a similar pattern, with 77 % fewer neurons than uninfected control mice at 6 wpi and 71% at 36 wpi. Benznidazole-mediated cure of infection led to recovery of neuron numbers by 36 wpi, with only 47% and 21% less than the uninfected control mean in the proximal and distal myenteric plexus respectively. Denervation in relapsed mice was significant, but of lower magnitude (69% proximal, 56% distal) than for infected-untreated mice (Fig. 2g, 2h), in line with their intermediate transit delay phenotype (Fig. 2b - f).

We observed a morphologically heterogeneous population of HuC/D^+^ myenteric neuronal bodies in colon samples from benznidazole-cured mice. A subset of these neurons resembled those seen in healthy control ganglia, while another subset appeared atypically smaller and rounder with weaker anti-HuC/D reactivity. These were commonly present in the same ganglion as neuronal cell bodies with normal morphology and neighbouring healthy control-like myenteric ganglia (Fig. 2g). Together, these data show that benznidazole treatment reverses the transit time and constipation phenotypes in the DCD mouse model, and that this is associated with substantial recovery of myenteric neuron density, particularly in the distal colon. In mice in which infection relapsed, transit time delay returns but not to levels observed in untreated infected controls, with an intermediate recovery of myenteric innervation.

### Distinct gene expression profiles for chronic, relapsed and cured infections

To further investigate how the balance of infection and host immunity impacts on the ENS during *T. cruzi* infection, we performed Nanostring multiplex analysis of host gene expression, focussing on immune response (n = 491) and ENS (n = 17) genes. In mice with untreated chronic infections (36 wpi), there were 128 significantly differentially expressed genes (DEGs) compared to uninfected controls in colon tissue, of which 108 were up- and 20 down-regulated (Fig. 3a, 3b). These evidenced a type 1-polarised inflammatory response involving class I and class II antigen presentation (e.g. *Ciita, Tap1, Psmb9, B2m; H2-Aa, H2-Ab1, Cd74*), cytokines/chemokines (e.g. *Ifng, Il21, Tnf; Cxcl9, Cxcl10, Ccl5*), transcription factors (e.g. *Stat1,-2,-3,-6, Irf1,-88*), cytotoxic lymphocyte markers and effectors (*Cd8a, Cd8b1, Cd226, Gzmb, Fasl, Prf1)*, complement factors (*C2, C6, C4a, C3, C7, C1qb*) and Fc receptors (*Fcgr1,-4,-3, Fcer1a,-1g*). There was also evidence of significant dysregulation at the pathway level for antigen presentation, interferon signalling, apoptosis and phagocytosis (Fig. 3d, Extended Data Fig. 3a). Consistent with the enduring capacity of *T. cruzi* to survive in this inflammatory environment and the need to prevent excessive GI tissue damage, the up-regulated DEG set included a diverse range of immuno-inhibitory mediators: *Btnl1, Btnl2, Cd274* (PD-L1), *Socs1, Lilrb4, Lilrb3, Lair1, Tnfaip3, Serping1*.

**Figure 3:**
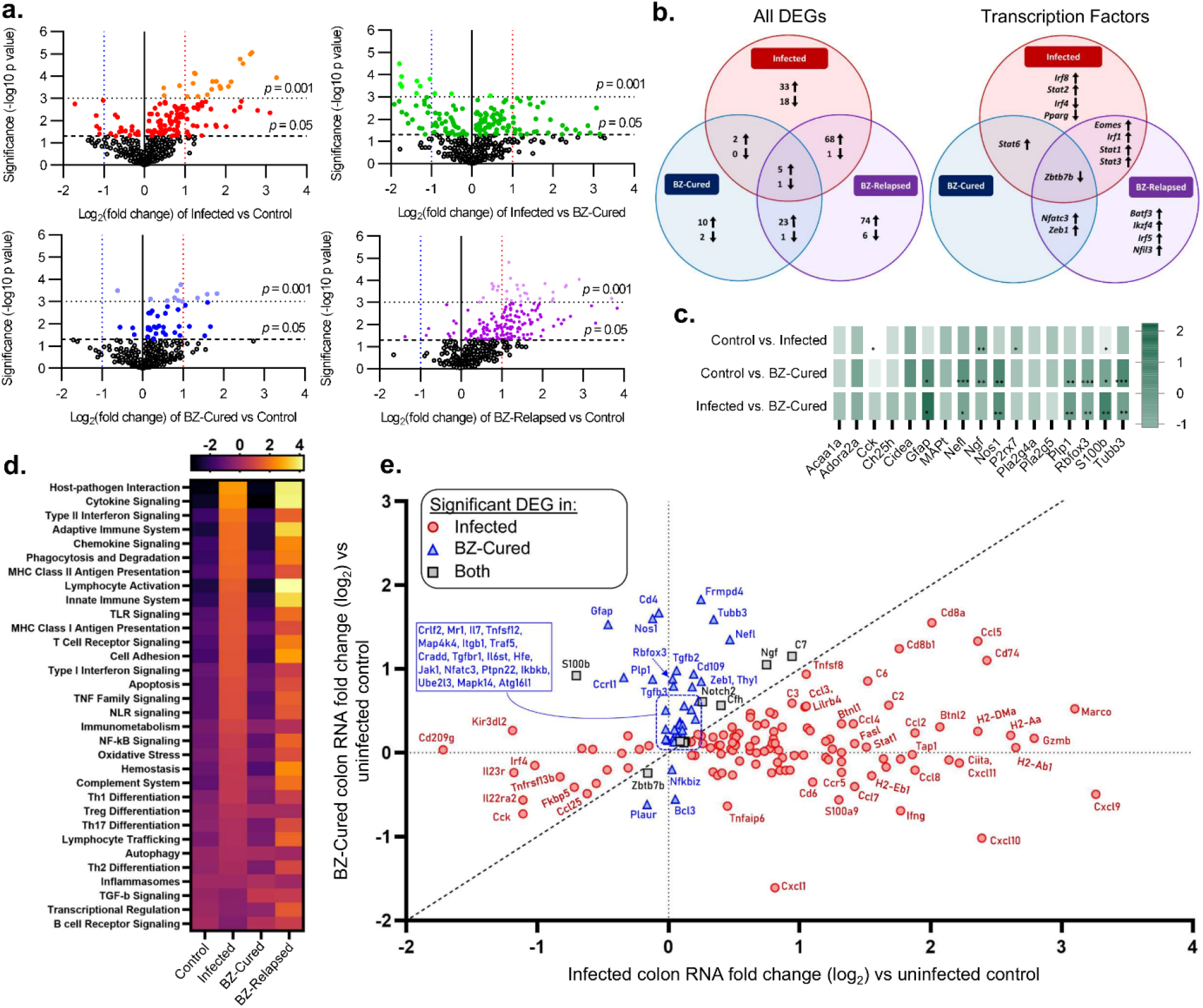
Enteric neuro-immune gene expression signatures associated with *T. cruzi* infection and benznidazole treatment outcomes. **a**, Volcano plots of the log_2_-transformed fold change and significance (−log_10_ *p* value) of differentially expressed genes (DEGs) in colon tissue from *T. cruzi* infected (red shades), benznidazole treated and cured (BZ-Cured; blue shades) and benznidazole treated and relapsed (BZ-Relapsed; purple shades) vs. uninfected control C3H/HeN mice; and BZ-Cured vs. infected (green shades) mice. Lighter coloured shaded dots indicate statistical significance of *P* < 0.05 and darker shaded dots *P* < 0.001. **b**, Venn diagrams show same-direction differentially expressing genes (DEG) shared between infected, BZ-Cured and BZ-Relapsed groups for all genes analysed (left, n shows number of genes) and transcription factors (right) (threshold of *p* < 0.05 vs uninfected controls in at least one of the groups). Arrows indicate up-and down-regulation. **c**, Relative change in neuro-glial genes between the indicated experimental groups; colour intensity corresponds to fold change (log_2_) expression vs uninfected controls. **d**, Signalling pathway scores for each group. **e**, Comparison of directionality and extent of gene expression change in chronically infected and BZ-cured mice vs uninfected controls (*n* = 163 genes that are significant DEGs in at least one group). Red circles are DEGs specific to the chronic, untreated infections, blue triangles are DEGs specific to the BZ-Cured mice and grey squares are DEGs shared by both groups. Diagonal dashed line is the line of equivalence. Vertical and horizontal dashed lines indicate position for genes with identical expression levels as uninfected controls in chronic infection and BZ-cured mice respectively. Infected and uninfected controls *n* = 6, BZ-Cured and BZ-Relapsed *n* = 3. Statistical significance was determined by 2-tailed, unpaired Student’s *t*-test for each gene (\**P* < 0.05, \*\**P* < 0.01, \*\*\**P* < 0.001, **** *P* < 0.0001).

Next, we evaluated the impact of benznidazole treatment on host gene expression in the contexts of sterile cure and infection relapse. Relapsed infections were associated with a larger DEG set (n = 179 vs uninfected) than untreated chronic infections. The majority, 58.6%, were same direction DEGs as chronic infections, but the data also revealed relapse-specific changes (Fig. 3b, Extended Data Fig. 4). Notably, these included stronger upregulation of 54 genes, including *Cd8a, Cd8b1, Ccl5, Cxcl1, Gzmb* and *Ifng*, and unique up-regulation of cytotoxic effectors *Gzma* and *Fas*, leukocyte markers *Cd4, Cd7* and *Cd27*, transcription factors (*Batf3, Ikzf4, Irf5, Nfil3*) and the components of integrins α4β1 (VLA-4), α5β1 (VLA-5), αLβ2 (LFA-1), αMβ2 (MAC-1). Given the lower and less disseminated parasite loads seen in these animals (Fig. 1), this broader gene expression profile is consistent with enhanced control of *T. cruzi*, enduring for months after non-curative treatment.

Colon tissue from benznidazole cured mice had gene transcript abundances equivalent to uninfected controls for 119 (93%) of the 128 DEGs that were identified in untreated chronic infections, indicating near complete reversion to homeostasis (Fig. 3). At the pathway level, there were no significant differences between cured and uninfected groups (Extended Data Fig. 4). Nevertheless, cured mice did have a distinct profile compared to uninfected control mice, comprising 44 DEGs (40 up-and 4 down-regulated genes) (Fig. 3b, 3c, 3e). Of note, the most up-regulated gene set was enriched for neuronal markers (*Rbfox3* (NeuN), *Nefl, Tubb3*), enteric glial cells (EGCs) (*S100b, Gfap, Plp1*), genes associated with neural development (*Ngf, Frmpd4*) and neurotransmission (*Nos1*). The EGC marker *S100b* was the only switched direction DEG, with reduced expression in chronic infections and increased expression in cured mice (Fig. 3e). Furthermore, multiple genes involved in tissue repair and regeneration were also highly significant DEGs in cured mice, including *Notch2, Tgfb2, Tgfbr1, Tgfb3* and *Zeb1*. Sterile cure of *T. cruzi* infection therefore results in dissipation of the chronic inflammatory environment in the colon and induction of a tissue repair program that encompasses key components of the ENS.

### Neuro-glial impact of benznidazole treatment of *T. cruzi* infections

Nitrergic neuronal inhibitory signalling is critical for homeostatic control of peristalsis and its disruption is linked to a range of enteric neuropathies, including human DCD (34). Evidence for altered *Nos1* gene expression in our experimental model ((33); Fig. 3c, 3e) led us to analyse the expression of the corresponding protein (neuronal NOS, nNOS) in myenteric neurons from the proximal and distal colon using immunofluorescence. The number of distal colon nNOS^+^ neurons was significantly reduced in untreated and relapse infections compared to uninfected controls (Fig. 4a, 4b). In benznidazole-cured animals, nNOS^+^ neuron density remained lower on average than uninfected controls, but the difference was not statistically significant (Fig. 4a, 4b).

**Figure 4:**
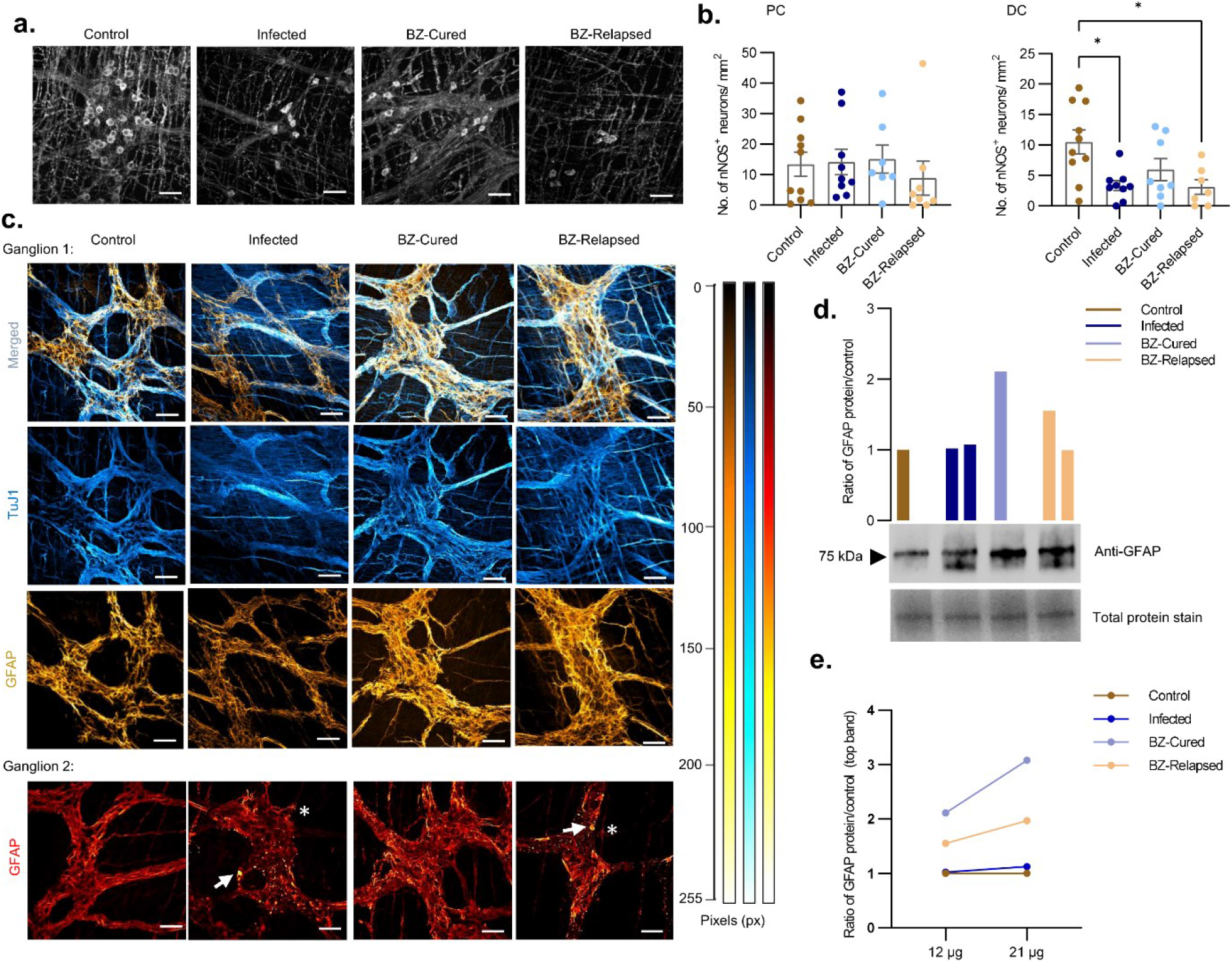
Enteric nitrergic neuron and glial cell dynamics in chronic *T. cruzi* infections and after benznidazole treatment. **a, b**, Representative immunofluorescence images (a) and quantification (b) of nNOS^+^ neurons in the colonic myenteric plexus of control (*n* = 10), infected (*n* = 9), benznidazole treated cured (BZ-Cured, *n* = 7 in proximal [PC] and *n* = 8 in distal colon [DC]) and benznidazole treated relapsed (BZ-Relapsed, *n* = 8 in PC and *n* = 7 in DC) mice. Bar plots show density of nNOS^+^ neuronal cell bodies. **c**, Representative immunofluorescence images of myenteric plexus enteric glial cells (EGCs, GFAP^+^, gold to yellow fluorescence intensity scale) and enteric neural network (TuJ1^+^, blue to cyan fluorescence intensity scale) in the indicated experimental groups. Top panel shows merged images of GFAP^+^ and TuJ1^+^ cells. Bottom panel shows images of morphologically diverse GFAP^+^ enteric glial cells in a second example ganglion revealed on a red to yellow fluorescence intensity scale. White arrows show aberrant EGC morphology and white stars show activated EGC morphology. All confocal images (**a** and **c**) were taken at 400X magnification, scale bar = 50 μm. Colour heat map scale shows pixel fluorescence intensity. **d, e**, Western blot analysis of GFAP in full thickness colon tissue lysates. Bar plot (d) and paired dot plot (e) show normalised GFAP protein abundance as a ratio of uninfected controls). For all groups, the sample analysed was a pooled lysate of 3 independent biological replicates (mice)). The blot in **d** shows α-GFAP staining using 12 μg of each lysate (corresponding to the bar plot groups above). Total protein was visualised using tryptophan trihalo-crosslinking gels for loading control and normalisation; the most abundant protein band is shown. GFAP abundance differences were validated by analysing replicate gels with two lysate quantities (12 and 21 μg) as indicated in plot **e**.

The RNA-based analyses strongly implicated EGCs in post-cure tissue repair (Fig. 3c, 3e, Extended Data Fig. 3b), so these were further analysed at the protein level. Immunofluorescence analysis of neuro-glial network morphology (Fig. 4c) clearly showed EGCs (glial fibrillary acidic protein, GFAP^+^) wrapped around neurons (neuron-specific β-tubulin, TuJ1^+^) in the myenteric plexus for all groups. Networks of GFAP^+^ EGCs in samples from untreated infections showed the weakest staining intensity and contained fragmented aggregates and patches in the plexus compared to controls, cured and relapsed mice. Conversely, GFAP staining in samples from benznidazole treated animals exhibited control-like glial morphology with a dense network, which was also preserved in relapsed infections (Fig. 4c). Western blot analysis of GFAP protein expression revealed bands at 75 kDa, larger than reference brain GFAP protein (50 kDa), which most likely reflects gut-specific post-translational modification. A second band was observed in both infected-untreated and relapse groups at approximately 70 kDa, indicating altered GFAP protein structure as a feature of DCD pathology. Benznidazole treatment induced doubling of the GFAP protein abundance compared to controls and disappearance of the secondary band, while the relapse infection group showed a more moderate increase overall and a minor secondary band (Fig. 4d, 4e). In combination, the RNA-and protein-based data provide evidence that sterile cure of *T. cruzi* infection is followed by increased EGC activity in the colonic myenteric plexus, which may contribute to the recovery of normal transit.

### Delayed treatment improves GI function to a lesser extent than early treatment

Most cases of human Chagas disease are associated with chronic *T. cruzi* infections. We therefore investigated treatment initiated at 24 wpi, to assess the impact on DCD in the chronic stage (Fig. 5). The bioluminescence profile of the infected-untreated mice followed a similar pattern as previously shown (Fig. 1a, 1e). Treatment with benznidazole at 24 wpi resulted in elimination of parasite bioluminescence by 30 wpi (Fig. 5a, 5b), a gradual gain of body weight (Extended Data Fig. 5a) and reversal of splenomegaly (Fig. 5c). Relapses of infection were detected in 30% (3/10) of the treated mice with reappearance of the bioluminescence signal, mostly in the abdominal area (Fig. 5a, 5b and 5d). *Ex vivo* imaging at end-point necropsy of infected mice (48 wpi) showed the highest intensity and frequency of infection in the heart and GI tract compared to other organs (Fig. 5e). Although there were only 3 relapse cases, the distribution of infections amongst organs and tissues appeared to have a similar profile to the acute stage treatment (Fig. 5e, 5f).

**Figure 5:**
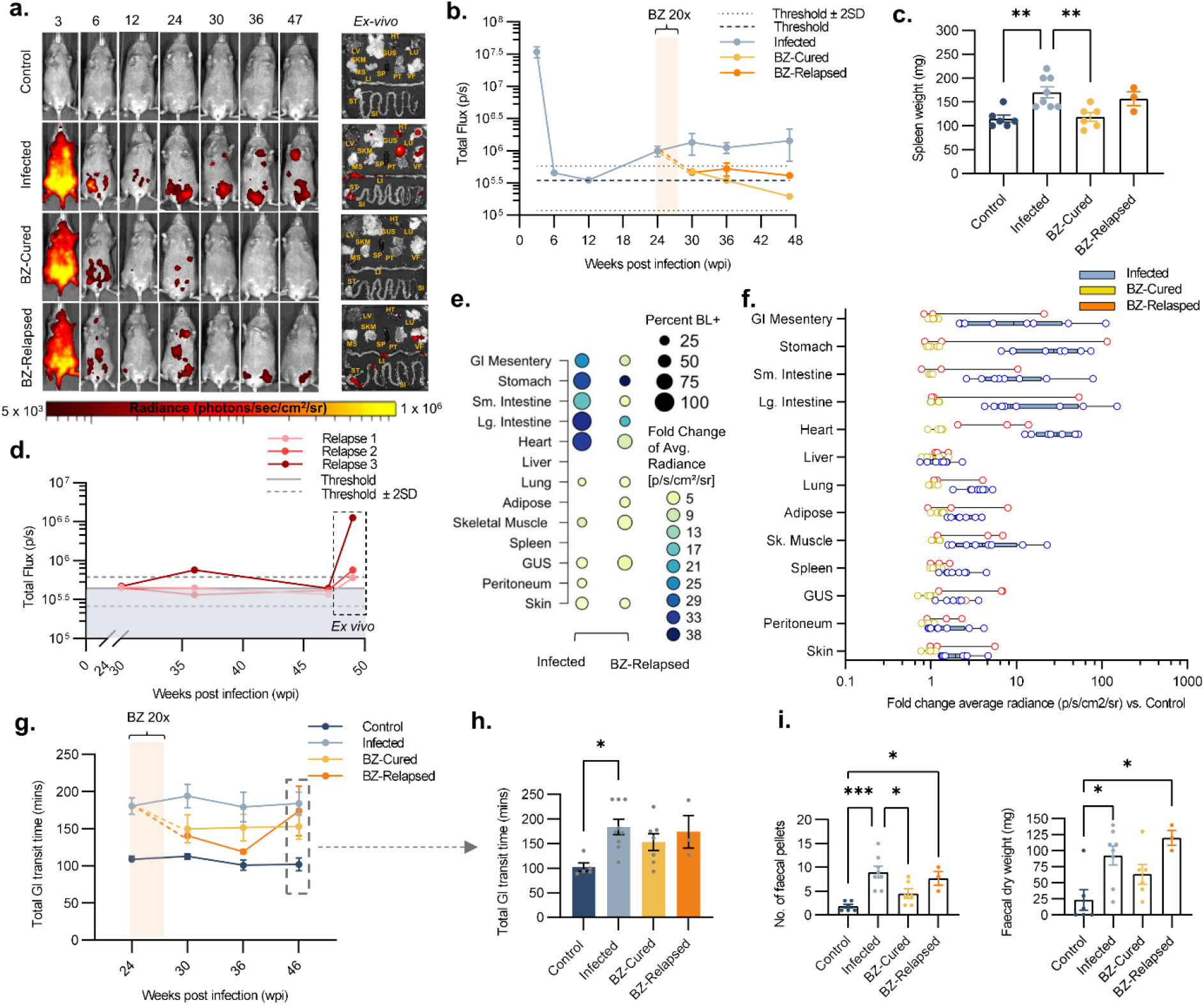
Impact of chronic phase benznidazole treatment on experimental digestive Chagas disease. **a, b** Representative *in vivo* bioluminescence (BL) image series (a) and quantification (b) of *T. cruzi*-infected C3H/HeN mice before and after treatment with benznidazole (BZ) at 24 weeks post-infection (wpi). *Ex vivo* images show distribution and intensity of infection in internal organs and tissue samples (liver - LV, lungs - LN, gut mesenteric tissue - MS, heart - HT, spleen - SP, skeletal muscle - SKM, visceral fat - VF, stomach - ST, small intestine - SI, large intestine - LI, genitourinary system - GUS, and peritoneum - PT) at necropsy. BL intensity (radiance) is expressed as log-scale pseudocolour heat maps. **I**nfected mice *n* = 8, except *n* = 19 at 6 and 12 wpi, *n* = 16 at 24 wpi and *n* = 20 at 3 wpi; benznidazole treated and cured (BZ-Cured; *n* = 7) and benznidazole treated and relapsed (BZ- Relapsed; *n* = 3). Dashed lines show thresholds: uninfected control mean auto-luminescence (dark blue dashed line) and mean ± 2SD (light blue dotted lines). **c**, Spleen weights of control (*n* = 6), infected (*n* = 8), BZ-Cured (*n* = 6) and BZ-Relapsed (*n* = 3) groups. **d**, *In vivo* BL intensity of individual BZ-Relapsed mice after treatment at 24 wpi. Thresholds shown as uninfected control mean auto-luminescence (grey line) and mean ± 2SD (grey dotted lines). **e**, Dot plot and **f**, box plot show relative *ex vivo* infection intensities in the indicated organs/tissue at 48 wpi. Data are expressed as the mean fold change in bioluminescent radiance vs. the uninfected control mean for groups (e) and individual mice (f). In dot plot **e**, the circle sizes indicate the percentage of mice with bioluminescence positive (BL^+^) signal and the circle colours show the infection intensity (fold change of mean bioluminescent radiance vs uninfected control). Infected *n* = 8, BZ-Cured *n* = 6 and BZ-Relapsed *n* = 3 mice. **g**, Mean GI transit time of control (*n* = 5, except *n* = 11 at 24 wpi and *n* = 6 at 30 wpi), infected (*n* = 9, except *n* = 23 at 24 wpi and *n* = 8 at 30 wpi), BZ-Cured (*n* = 7) and BZ-Relapsed (*n* = 3) mice. Cream overlaid bar shows BZ treatment window (24 - 27 wpi, 20 daily doses). **h**, Bar plots show final GI transit times at 46 wpi for individual mice. Control *n* = 5, infected *n* = 9, BZ-Cured *n* = 7 and BZ-Relapsed *n* = 3. **i**, Bar plots show number of faecal pellets number (left) and dry weight (right) retained in the colon after 4 hours of fasting at 48 wpi for control (*n* = 6), infected (*n* = 8), BZ-Cured (*n* = 6) and BZ-Relapsed (*n* = 3) mice. Data are expressed as mean ± SEM. Statistical significance was tested using one-way ANOVA followed by Tukey’s HSD test (\**P* < 0.05, **** *P* < 0.0001).

As expected, there was a significantly longer GI transit time (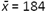 mins) in infected, untreated mice at all chronic time-points compared to uninfected controls (Fig. 5g, 5h). Transit time in sterile cured mice improved to an intermediate level (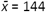 mins), which was stable over for the duration of the experiment, indicating a partial recovery of function (Fig. 5g, 5h). The relapse mice initially showed strong improvements in transit time post-treatment, but by the end point, they transitioned towards an increased transit time (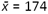 mins), close to the delay seen in infected-untreated mice (Fig. 5g, 5h). When we analysed colonic faecal retention, there was stronger evidence for recovery of GI function in cured mice. This constipation phenotype was significantly alleviated after benznidazole-mediated cure of infection, while there was no significant improvement in mice where infection relapsed after treatment (Fig. 5i and Extended Data Fig. 5e). Retrospective comparison of pre-treatment body weights, parasite loads and transit times showed that there were no significant differences between benznidazole treated mice that were cured compared to relapsed (Extended Data Fig. 6). In summary, when treatment was delayed until the chronic phase of infection, the observed sterile parasitological cure rate was slightly higher than for treatment at 6 weeks (70% vs 59%) and there was more modest recovery of GI transit function. Therefore, the timing of anti-parasitic treatment is an important factor affecting the degree to which GI function can be restored in DCD.

## Discussion

Treatment for DCD is limited to palliative dietary adjustments and surgical interventions with significant mortality risk (16). Lack of data has prevented a consensus on whether *T. cruzi* infected adults who are asymptomatic should be treated with anti-parasitic chemotherapy (4, 12, 35). Our findings in a pre-clinical mouse model provide evidence that prompt treatment with benznidazole can prevent chronic DCD. When treatment was delayed until the chronic phase, sterile cure of infection only resulted in a partial GI functional recovery, indicating that some tissue pathology reaches an irreversible stage. This echoes the results of clinical trials of benznidazole in late-stage chronic Chagas cardiomyopathy patients, in which the drug performed no better than placebo in preventing disease progression or death (8). Further work is required to determine the point at which cure of infection will cease to yield functional improvement in DCD. Nonetheless, our findings provide a pre-clinical *in vivo* evidence base supporting the concept that the earlier anti-parasitic chemotherapy is initiated, the greater the chances of preventing or delaying the progression of digestive disease.

In the context of DCD, our results provide insight into the dynamic relationships between *T. cruzi* infection, host responses, ENS damage and tissue repair. Most denervation occurred in the acute phase of infection, yet the complete and sustained normalisation of GI transit function after benznidazole treatment at 6 weeks shows that these acute losses are not sufficient to explain chronic disease symptoms, contrary to early theories of DCD aetiology (18). Transient functional improvement in untreated controls in the early chronic phase further supports this conclusion (33). Over time, chronic infection of the GI tract led to further neuron losses and gradual decline in GI function.Moreover, in cases of failed treatment, relapses of *T. cruzi* infection were associated with a return to GI dysfunction. Together, these data support the conclusion that chronic infection actively drives disease, as has been circumstantially suspected from the detection of parasites in GI tissues from human DCD patients (19-27). However, the disease aetiology is more complex than anticipated because neither parasite load, nor the degree of denervation directly predicted the severity of functional impairment. This might be explained by compensatory plasticity of the remaining ENS and/or extrinsic circuitry in the denervated colon (36). As seen in the CNS, it is also possible that the existing ENS neural circuit rewires to compensate for the neuronal loss by rebalancing the excitatory and inhibitory outputs of the network (37). Our neuro-immune gene expression analysis indicated that the broader balance of inflammatory, regulatory and tissue repair factors at play in the infected colon also contributes to the DCD phenotype.

Why the GI tract serves as a long-term permissive site for *T. cruzi* has been unclear because there have been minimal data on gut-specific immune responses. At the transcriptional level, we found that chronic infection of the colon is associated with a robust type 1 inflammatory response, dominated by markers of CD8^+^ T cell recruitment, similar to that seen in studies of other tissue types (10, 38-42). The discovery that at least 9 genes with immuno-inhibitory potential were also upregulated suggests that there are host-intrinsic mechanisms that limit tissue damage, yet also enable *T. cruzi* persistence. We previously found that chronic *T. cruzi* infection is restricted to a few rare foci, which can be widely scattered within the colonic smooth muscle layers and are regularly re-seeded in new locations by motile trypomastigotes (33, 43). Here, we analysed RNA from full thickness colon tissue and, for selected markers of neurons and glia, immuno-fluorescence analysis of the myenteric plexus. Therefore, a key challenge now will be to determine how infection and immune response dynamics relate to specific cell types, and how these in turn shape pathology at finer spatial and temporal scales.

The extent to which the adult ENS is capable of repair and regeneration is a fundamental question, with relevance across diverse enteric neuropathies. We observed a recovery of myenteric plexus neuron numbers in mice several months after sterile cure of *T. cruzi* infection. There are several potential neurogenic mechanisms that could underly this replenishment. It has been proposed that enteric neurons are regularly replaced from a population of neural stem cells as part of gut homeostasis (44), however, the majority of studies indicate that enteric neurogenesis is absent, or extremely limited in the steady-state adult gut (45-50). Regeneration of the ENS involves neurogenesis from enteric glial cell pre-cursors after chemical injury using benzalkonium chloride (BAC) or Dextran Sulfate Sodium (DSS) (46, 47, 49, 50) and from extrinsic Schwann cell pre-cursors in mouse models of Hirschprung’s disease (51, 52). Our approach did not enable us to define the ontogeny of new neurons, but the data suggest that glial cells are likely to be involved. We saw upregulation of the canonical glial markers GFAP and S100β specifically in sterile cured mice, as well as PLP1, which corresponds to the subset of glial cells that differentiate into neurons in the DSS colitis model (47). Transcriptional changes in healing colon tissue also comprised up-regulation of the stem cell transcription factor *Zeb1*, which regulates epithelial–mesenchymal transition (53) and neuronal differentiation in the CNS (54), including from radial glial-like cells in the adult hippocampus (55). We also saw up-regulation of *Ngf* and *Notch2*, critical regulators of neurogenesis; Notch signalling in particular has been implicated in maintenance of the neural progenitor pool and controlling glial to neuronal differentiation in the CNS (56) and the gut (57, 58). The *T. cruzi* infection and cure approach may therefore open new experimental opportunities to study adult neurogenesis and inform the development of regenerative therapies for enteric neuropathies, most especially DCD.

## Materials and Methods

### Parasites

This study used the TcI-JR strain of *T. cruzi* parasites constitutively expressing the red-shifted firefly luciferase variant *PPy*RE9h, as previously described (29). Epimastigotes were cultured *in vitro* in supplemented RPMI-1640 medium at 28°C under selective drug pressure with 150 μg ml^-1^ G418. MA104 monkey kidney epithelial cell monolayers were infected with metacyclic trypomastigotes, obtained from stationary phase cultures, in MEM media + 5% FBS at 37°C and 5% CO_2_. After 5 to 10 days, tissue culture parasites (TCTs) were harvested from the culture supernatant and aliquots from a single batch were cryopreserved in 10% DMSO. For *in vivo* infections, TCTs were thawed at room temperature, sedimented by centrifugation at 10,000x *g* for 5 minutes, washed in 1 ml complete medium, sedimented again and resuspended in 250 μl complete medium. After 1 hr incubation at 37°C, active parasites were counted and the suspension was adjusted to the required density.

### Animals and Infections

All animal procedures were performed under UK Home Office project license P9AEE04E, approved by LSHTM Animal Welfare Ethical Review Board and in accordance with Animal Scientific Procedure Act (ASPA) 1986 regulations. Female C3H/HeN mice, aged 6-8 weeks, were purchased from Charles River (UK) and habituated for 1-2 weeks before experiments. All CB17 SCID mice used were female and bred in-house. Mice were housed in individually ventilated cages on a 12 hr light/dark cycle. They had access to food and water available *ad libitum* unless otherwise stated. Mice were maintained under specific pathogen-free conditions. Humane end-points were loss of >20% body weight, reluctance to feed or drink freely for more than 4-6 hours, loss of balance or immobility.

Inocula of 1 × 10^5^ TCTs were used to infect SCID mice via i.p. injection. After 3 weeks, motile blood trypomastigotes (BTs) were derived from the supernatant of cardiac whole blood after passive sedimentation of mouse cells for 1 hr at 37°C. C3H/HeN mice were infected with 1 × 10^3^ BTs in 0.2 ml PBS via i.p. injection. The benznidazole treatment schedule was 100 mg kg^-1^ day^-1^ for 20 consecutive days via oral gavage. Benznidazole was prepared from powder form at 10 mg ml^-1^ by dissolving in vehicle solution (0.5 % w/v hydroxypropyl methylcellulose, 0.5% v/v benzyl alcohol, 0.4% v/v Tween 80 in deionised water).

At experimental end-points, mice were killed by exsanguination under terminal anaesthesia (Euthatal/Dolethal 60 mg kg^−1^, i.p.) or by cervical dislocation. Selected organs and tissue samples were cleaned with PBS and either snap-frozen on dry ice, fixed in 10 % Glyofixx or transferred to ice-cold DMEM medium to suit different downstream analysis methods.

### Total GI transit time assay

Carmine red dye solution, 6% in 0.5% methyl cellulose (w/v) in distilled water was administered to mice by oral gavage (200 μl). Mice were returned to their home cage for 75 mins, after which they were placed in individual containers and observed. The time of excretion of the first red-stained faecal pellet was recorded and the mouse was returned to its cage. A cut off time of 4 hours was employed as the maximum GI transit delay for the assay for welfare reasons. Total GI transit time was calculated as the time taken to expel the first red pellet from the time of gavage.

### *In vivo* bioluminescence imaging

Mice were injected with 150 mg kg^−1^ D-luciferin i.p., then anaesthetised using 2.5% (v/v) gaseous isoflurane in oxygen. Bioluminescence imaging was performed after 10-20 minutes using an IVIS Lumina II or Spectrum system (PerkinElmer). Image acquisition settings were adjusted dependent on signal saturation (exposure time: 1 – 5 min; binning: medium to large). After imaging, mice were placed on a heat pad for revival and returned to cages. Whole body regions of interest (ROIs) were drawn on acquired images to quantify bioluminescence, expressed as total flux (photons sec^-1^), to estimate *in vivo* parasite burden in live mice (28). The detection threshold was determined using uninfected control mice. All bioluminescence data were analysed using Living Image v4.7.3.

### *Ex vivo* bioluminescence imaging

Food was withdrawn from cages 4 hours prior to euthanasia. Five to seven minutes prior to euthanasia, mice were injected with 150 mg kg^-1^ D-luciferin i.p.. After euthanasia, mice were perfused trans-cardially with 10 ml of 0.3 mg ml^-1^ D-luciferin in PBS. Typically, organs collected included heart, liver, spleen, lungs, skin, peritoneum, the GI tract, the genitourinary system and their associated mesenteries, as well as samples of hindlimb skeletal muscle and visceral adipose. These were soaked in PBS containing 0.3 mg ml^-1^ D-luciferin prior to imaging, which was performed as described above. Parasite load in each organ tissue was quantified using a measure of infection intensity. To do this, bioluminescence per organ or tissue sample was calculated by outlining ROIs on each sample and expressed as radiance (photons sec^−1^ cm^−2^ sr^−1^). Radiances from equivalent organs/tissues of age-matched, uninfected control mice were measured and the fold change in bioluminescence for each test sample was calculated. The detection threshold was determined by scoring the presence of *T. cruzi* infection foci of at least 10 contiguous bioluminescent pixels at radiance ≥3 × 10^3^ photons sec^−1^ cm^−2^ sr^−1^. These criteria were established by reference to a set of images from uninfected control animals (59).

### Faecal analyses

Isolated colon tissue was cleaned externally with PBS and faecal pellets were gently teased out of the lumen. Faecal pellets were counted and collected in 1.5 ml tubes. Wet weights were recorded and then the tubes were left to dry in a laminar flow cabinet overnight; dry weights were measured the following day.

### Histopathology

Paraffin-embedded fixed tissue blocks were prepared and 3-5 μm sections were stained with haematoxylin and eosin as described (29). Images were acquired using a Leica DFC295 camera attached to a Leica DM3000 microscope. For analysis of inflammation, nuclei were counted automatically using the Leica Application Suite V4.5 software (Leica).

### Immunofluorescence analysis

After necropsy, excised colon tissues were transferred from ice-cold DMEM to PBS. Tissues were cut open along the mesentery line, rinsed with PBS, then stretched and pinned on Sylgard 184 plates. Under a dissection microscope, the mucosal layer was carefully peeled away using forceps and the remaining muscularis wall tissue was fixed in paraformaldehyde (4% w/v in PBS) for 45 minutes at room temperature. Next, tissues were washed with PBS for 45 minutes, with 3 changes, at room temperature and permeabilised with PBS containing 0.5% Triton X-100 for 2 hours, followed by blocking for 1 hour (10% sheep serum in PBS containing 0.5% Triton X-100). Tissues were incubated with primary antibodies (mouse anti-HuC/D IgG clone 16A11 at 1:200 [ThermoFisher], rabbit anti-tubulin β-3 (TuJ1) polyclonal IgG at 1:500 [Biolegend], rat anti-GFAP monoclonal IgG clone 2.2B10 at 1:500 [ThermoFisher], rabbit anti-nNos polyclonal IgG at 1:500 [ThermoFisher]) in PBS containing 0.5% Triton X-100 for 48 hours at 4°C. Tissues were washed with PBS for 30 minutes with three changes, then incubated with secondary IgG (goat anti-mouse Alexa546, goat anti-rabbit Alexa633, goat anti-rat, all 1:500, ThermoFisher) in PBS containing 0.5% Triton X-100 for 2 hours and counterstained with Hoechst 33342 (1 μg ml^-1^) at room temperature. Control tissues were incubated with only secondary antibodies (without primary antibodies) to assess antibody specificity. Tissues were mounted on glass slides using FluorSave mounting medium (Merck).

Whole mounts were examined and imaged with a LSM880 confocal microscope using a 40X objective (Zeiss, Germany). Images were captured as Z-stack scans of 21 digital slices with interval of 1 μm optical thickness. Five Z-stacks were acquired per region (proximal and distal colon), per animal. Cell counts were performed on Z-stacks after compression into a composite image using the cell counter plug-in of FIJI software. Neuronal density was calculated as the number of HuC/D^+^ or nNOS^+^ neuron cell bodies per mm^2^. HuC/D signal was associated with high background outside ganglia in samples from infected mice, attributed to binding of the secondary anti-mouse IgG to endogenous IgG, so ENS-specific analysis was aided by anti-TuJ1 co-labelling and assessment of soma morphology.

### Western Blot

Frozen colon tissue samples were lysed in RIPA Buffer and the total protein concentration was quantified using a BCA assay kit as per manufacturer’s protocol (ThermoFisher). Lysates were obtained from three independent biological samples per group and pooled into a single sample for analysis. Polyacrylamide gel electrophoresis was performed to separate proteins using 4-20% stain-free TGX gels (Bio-Rad). Proteins were visualised by UV-induced fluorescence using a Chemidoc imaging system (Bio-Rad) to verify equal loading of samples. The most abundant protein band in each loading control sample was used for quantification. Proteins were transferred to nitrocellulose membranes in a trans-blot turbo transfer system (Bio-Rad). Membranes were blocked for 30 minutes using 5% skimmed milk in PBST and then probed with rat anti-GFAP primary antibody (1:2000, cat # 13-0300, Thermo Scientific) overnight at 4°C, followed by incubation with HRP-conjugated goat anti-rat secondary antibody (1:5000, cat # 31460 Thermo Scientific) for 2 hours at room temperature and visualisation using enhanced chemiluminescence (ECL kit, GE Healthcare Life Sciences). Data were analysed using the gel analysis package in FIJI.

### RNA extraction

Frozen tissue samples were thawed in 1 ml Trizol (Invitrogen) per 30-50 mg tissue and immediately homogenised using a Precellys 24 homogeniser (Bertin). 200 μl of chloroform was added to each sample and mixed by vortex. The aqueous phase was separated by centrifugation at 13,000 g at 4°C and RNA was purified using the RNeasy Mini Kit (Qiagen) with on-column DNAse digestion, as per manufacturer’s protocol. A Qubit Fluorimeter (ThermoFisher) and/or a Nanodrop instrument was used to assess RNA quality and quantity.

### RT-qPCR

cDNA was synthesised from 1 μg of total RNA using Superscript IV VILO mastermix (Invitrogen), as per manufacturer’s protocol, in reaction volumes of 20 μl. qPCR reactions were carried out using QuantiTect SYBR green master mix (Qiagen) with 200 nM of each primer and 4 μl of cDNA diluted 1/50 in DEPC water. Reactions were run using an Applied Biosystems Fast 7500 machine (ThermoFisher) as per manufacturer’s protocol.

A final cDNA volume of 100 μl was made by adding RNase-free DEPC water (1: 5 dilution) and stored at −20 °C until further use. qPCR reactions consisted of 10μl QuantiTect SYBR green master mix, 6 μl of forward and reverse primer mix (200 nM; design in Supplementary Table 1) and 4 μl of cDNA diluted 1/50 in DEPC water. For No-RT and no template control reactions, 4 μl of solution from the No-RT cDNA reaction and DEPC water were added respectively. Reactions were run using an Applied Biosystems Fast 7500 machine (ThermoFisher) as per manufacturer’s protocol. Data were analysed by the ΔΔCt method (60) using murine *Oaz1* as the endogenous control gene.

### Nanostring gene expression analysis

RNA was adjusted to 30-60 ng μl^-1^ and analysed on a Nanostring nCounter system (Newcastle University, UK). We used a “PanelPlus” set of target probes comprising the standard mouse immunology nCounter codeset (XT-CSO-MIM1-12) and a custom selection of 20 probes from the mouse neuroinflammation and neuropathology codesets: *Acaa1a, Adora2a, Cck, Ch25h, Cidea, Drd1, Drd2, Gfap, MAPt, Nefl, Ngf, Nos1, Npy, P2rx7, Pla2g4a, Pla2g5, Plp1, Rbfox3, S100b* and *Tubb3*. The core immunology codeset comprised 547 protein-coding test genes, 14 house-keeping control genes, 6 positive binding control probes and 8 negative binding control probes. Fifty nine test genes were below a detection threshold limit (mean negative control bound probe count + 3 SDs) and were excluded from the analysis. The final codeset comprised probes for 508 test genes and was analysed using nSolver 4.0. Data were normalised in the Basic Analysis module with positive control and housekeeping gene normalisation probe parameters both set to geometric mean. Normalised data were then imported to the Advanced Analysis module and used to analyse differential gene expression between groups and pathway scores using default parameters. Samples were annotated with their run number as a confounding variable.

### Statistics

Individual animals were used as the unit of analysis. No blinding or randomisation protocols were used. Statistical differences between groups were evaluated using 2-tailed, unpaired Student’s *t*-test one-way ANOVA with Tukey’s post-hoc correction for multiple comparisons. These tests were performed in nSolver 4.0, GraphPad Prism v.8 or R v3.6.3. Differences of *p* < 0.05 were considered significant.

## Acknowledgements

We thank Hernán Carrasco for sharing parasite strains, Jody Phelan for helping with R scripts and the LSHTM Biological Services Facility staff for technical support and animal husbandry. The work was funded by an MRC New Investigator Research Grant (MR/R021430/1) and an EU Marie Curie Fellowship (grant agreement no. 625810).

## Extended Data

**Extended Data Figure 1:**
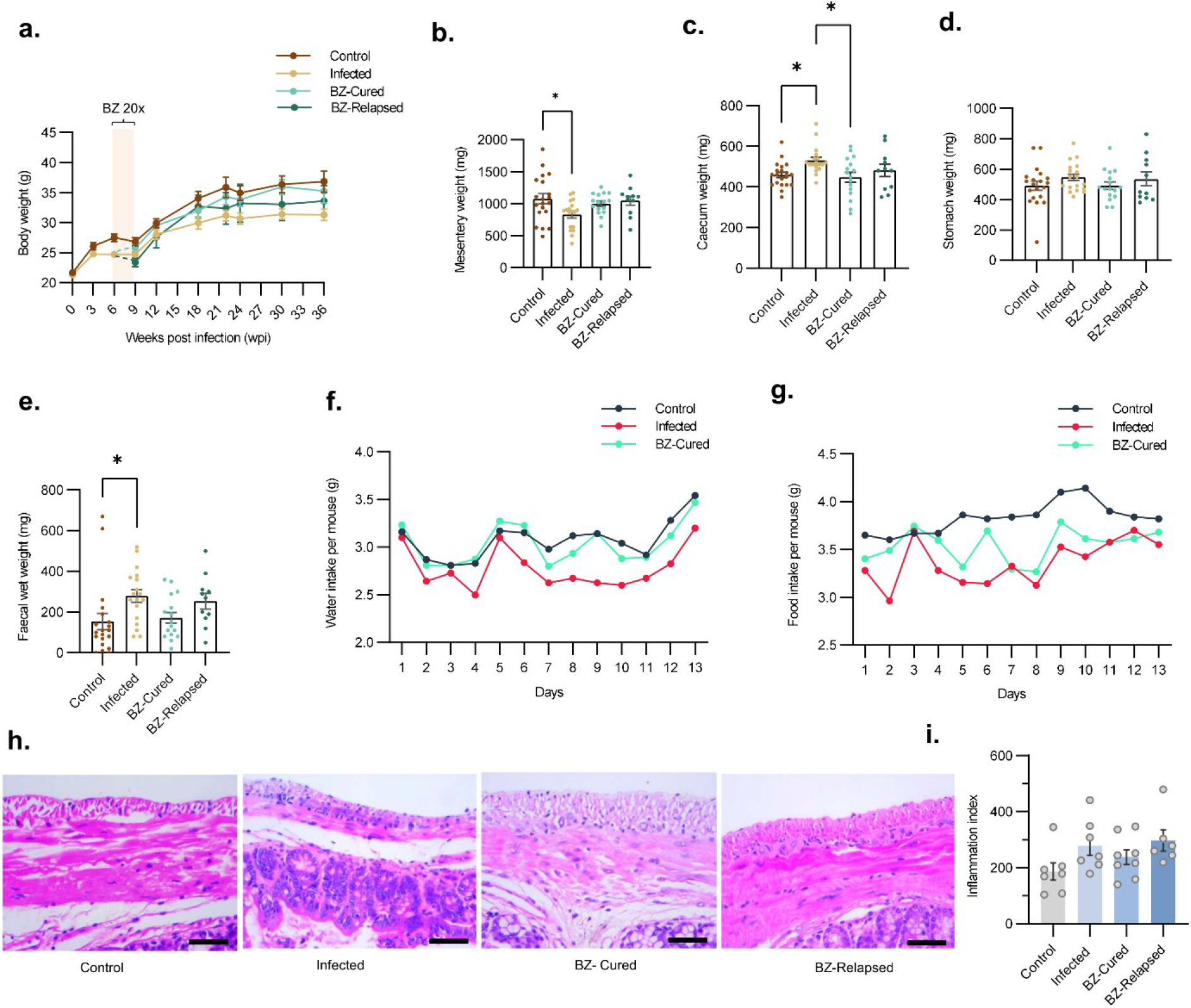
Additional GI assays after benznidazole treatment initiated at 6 weeks post-infection. **a**, Body weight line plots of control (*n* = 15; except *n* = 10 at 3, 6, 9, 22 and 36 wpi), infected (*n* = 10, except *n* = 25 at 0 and 3 wpi, and *n* = 24 at 6 wpi), benznidazole treated and cured (BZ-Cured; *n* = 7, except *n* = 8 at 9, 12 and 18 wpi) and benznidazole treated and relapsed (BZ-Relapsed; *n* = 5) mice. Cream bar on line plots show benznidazole treatment window (6 - 9 wpi). Bar plots show **b**, GI mesentery **c**, caecum **d**, stomach and **e**, wet faecal pellet weight of control (*n* = 20), infected (*n* = 18), BZ-Cured (*n* = 16) and BZ-Relapsed (*n* = 11) mice. **f**, Average daily water and, **g**, food intake per mouse of uninfected control (*n* = 2 cages, 10 mice), infected (*n* = 4 cages, 20 mice) and BZ-treated (*n* = 6 cages, 30 mice) mice over two weeks (35 – 37 wpi). **h**, Representative brightfield images of 5 μm thick colon transverse sections stained with haematoxylin-eosin (mucosa bottom; smooth muscle layers top). Images were taken at 400X magnification, scale bar at 50 μm. **i**, Adjacent bar plot shows number of nuclei per field to quantify cellular infiltration in control (*n* = 7), infected (*n* = 7), BZ-Cured (*n* = 8) and BZ-Relapsed (*n* = 6) mice. End-point data **b, c, d, e, h** and **i** are from 36 wpi. Statistical significance was tested using one-way ANOVA followed by Tukey’s HSD test (\**P* < 0.05).

**Extended Data Figure 2:**
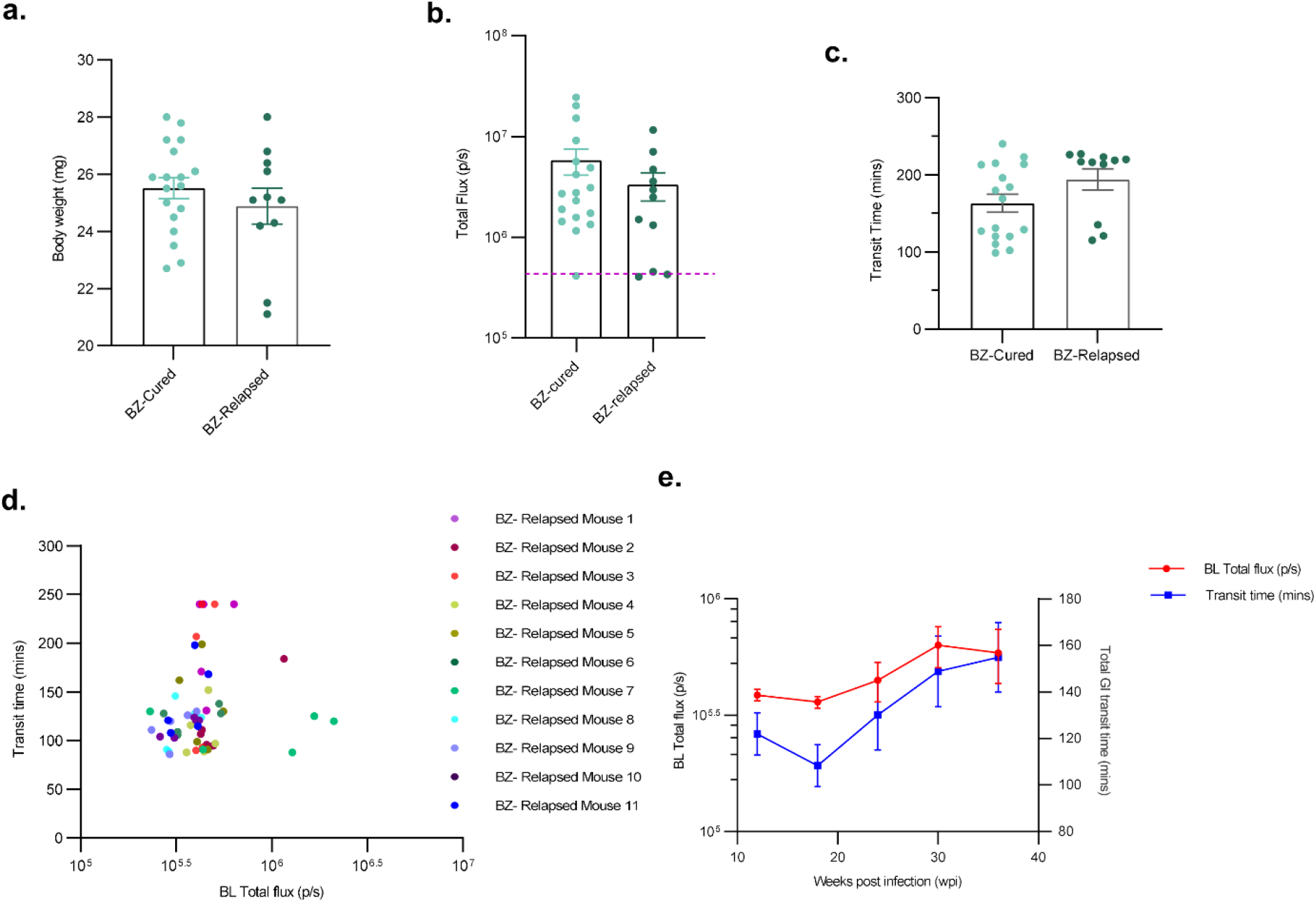
Pre-treatment characteristics and relapse kinetics after benznidazole treatment initiated at 6 weeks post-infection. **a-c**, Bar plots show characteristics of BZ-Cured (*n* = 18) and BZ-Relapsed (*n* = 11) mice retrospectively segregated at the point of treatment initiation (6 weeks post-infection (wpi)): **a**, body weight; **b**, bioluminescence intensity (total flux); and **c**, GI transit time. **d, e**, Comparison of parasite load (expressed as total flux) and transit time in BZ-Relapsed (*n* = 11) mice. The dot plot (d) shows pairwise correlation for all time points for individual animals, while the line plot (e) shows the group mean (+/-SEM) for bioluminescence (red line, left y axis) and transit time (blue line, right y axis) over time. Experimental end-point was at 36 wpi. Statistical significance was tested using one-way ANOVA followed by Tukey’s HSD test (\**P* < 0.05, \*\**P* < 0.01, \*\*\**P* < 0.001, **** *P* < 0.0001).

**Extended Data Figure 3:**
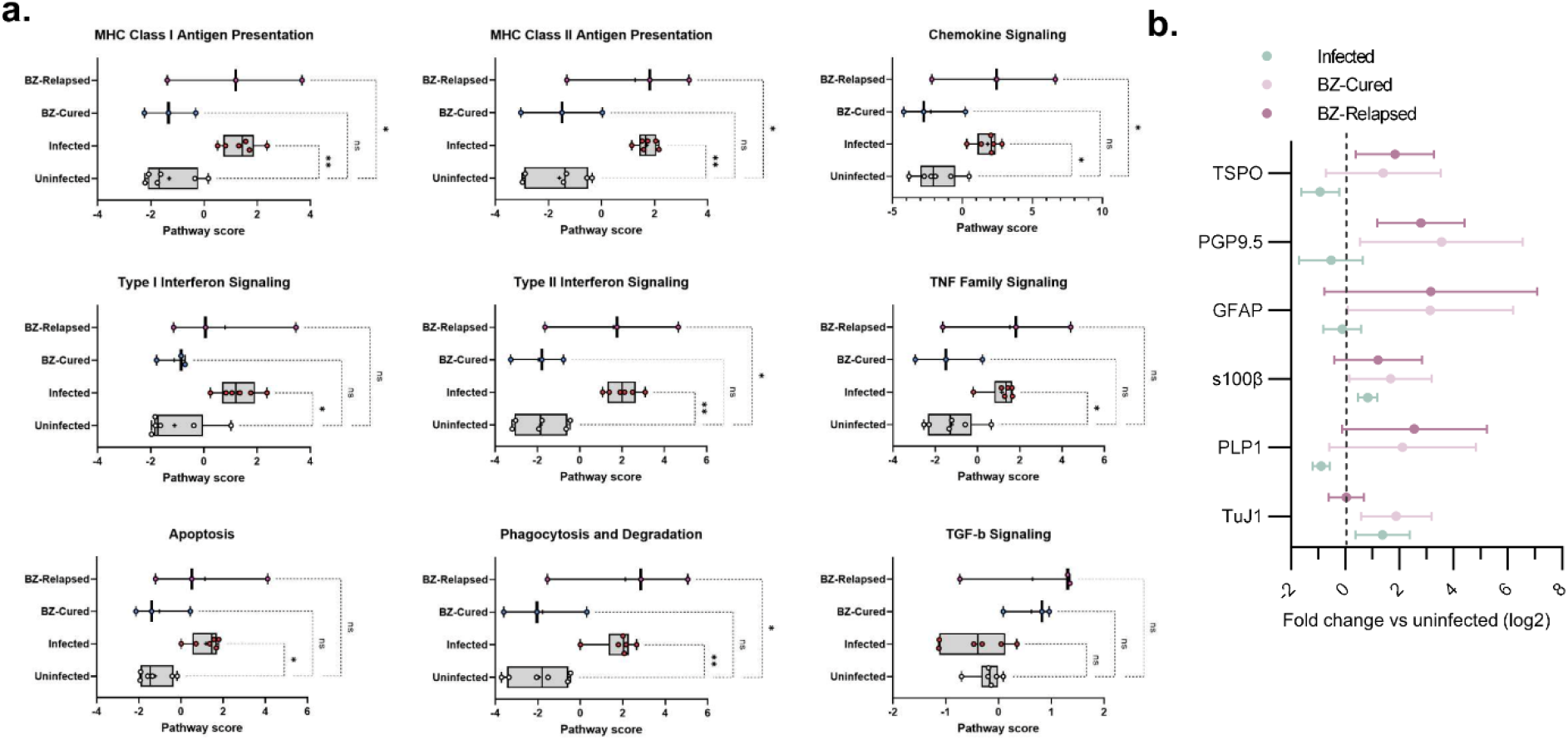
Gene expression analysis of immunoregulatory and neuro-glial genes. **a**, Pathway level analysis of immunoregulatory differential gene expression (*n* = 4 all) and **b**, qPCR analysis of additional neuronal and glial cell marker gene expression (*n* = 4 all). All mice were treated with benznidazole at 6 weeks post-infection (wpi) and RNA was extracted from colon tissue harvested at 36 wpi.

**Extended Data Figure 4:**
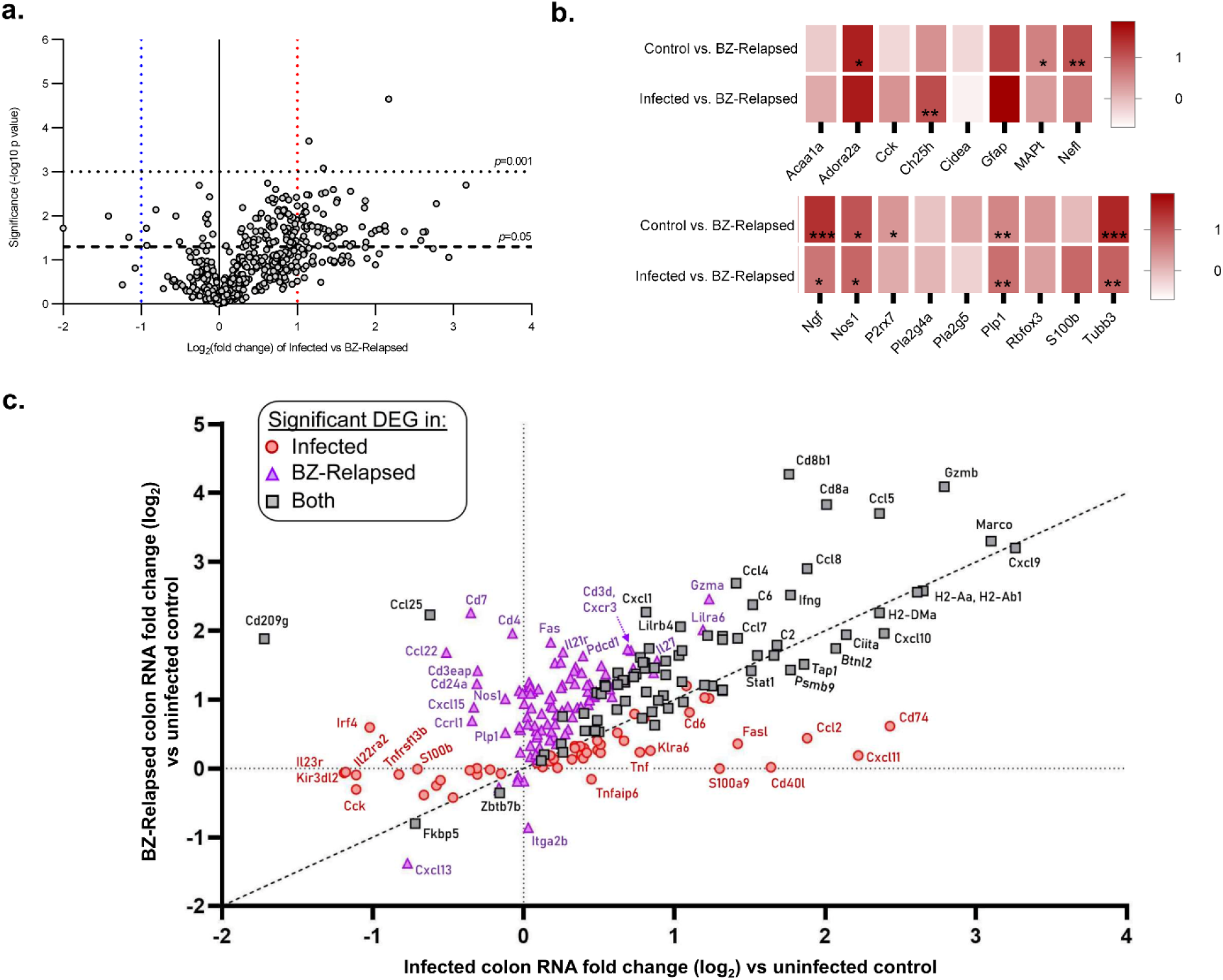
Analysis of differential gene expression in post-treatment relapsed infections. **a**, Volcano plot of the log_2_-transformed fold change and significance (−log_10_ *p* value) of differentially expressed genes (DEGs) in colon tissue from infected vs. benznidazole-treated relapsed (BZ-Relapsed) mice. **b**, Heat maps show relative change in neuro-glial genes between BZ-Relapsed vs. control (uninfected) or infected groups. Colour intensity indicates fold change (log_2_) expression level. **c**, Comparison of directionality and extent of gene expression change in infected and BZ-Relapsed mice vs controls (*n* = 230 genes that are significant DEGs in at least one group). Red circles are DEGs specific to the infected group, purple triangles are DEGs specific to the BZ-Cured mice and grey squares are DEGs shared by both groups. Diagonal dashed line is the line of equivalence. Vertical and horizontal dashed lines indicate position for genes with identical expression levels as controls in infected and BZ-Relapsed mice respectively. Infected and controls *n* = 6, BZ-Relapsed *n* = 3. All mice were treated with benznidazole at 6 weeks post-infection and RNA was extracted from colon tissue harvested at 36 wpi. Statistical significance was determined by 2-tailed, unpaired Student’s *t*-test for each gene (\**P* < 0.05, \*\**P* < 0.01, \*\*\**P* < 0.001, **** *P* < 0.0001).

**Extended Data Figure 5:**
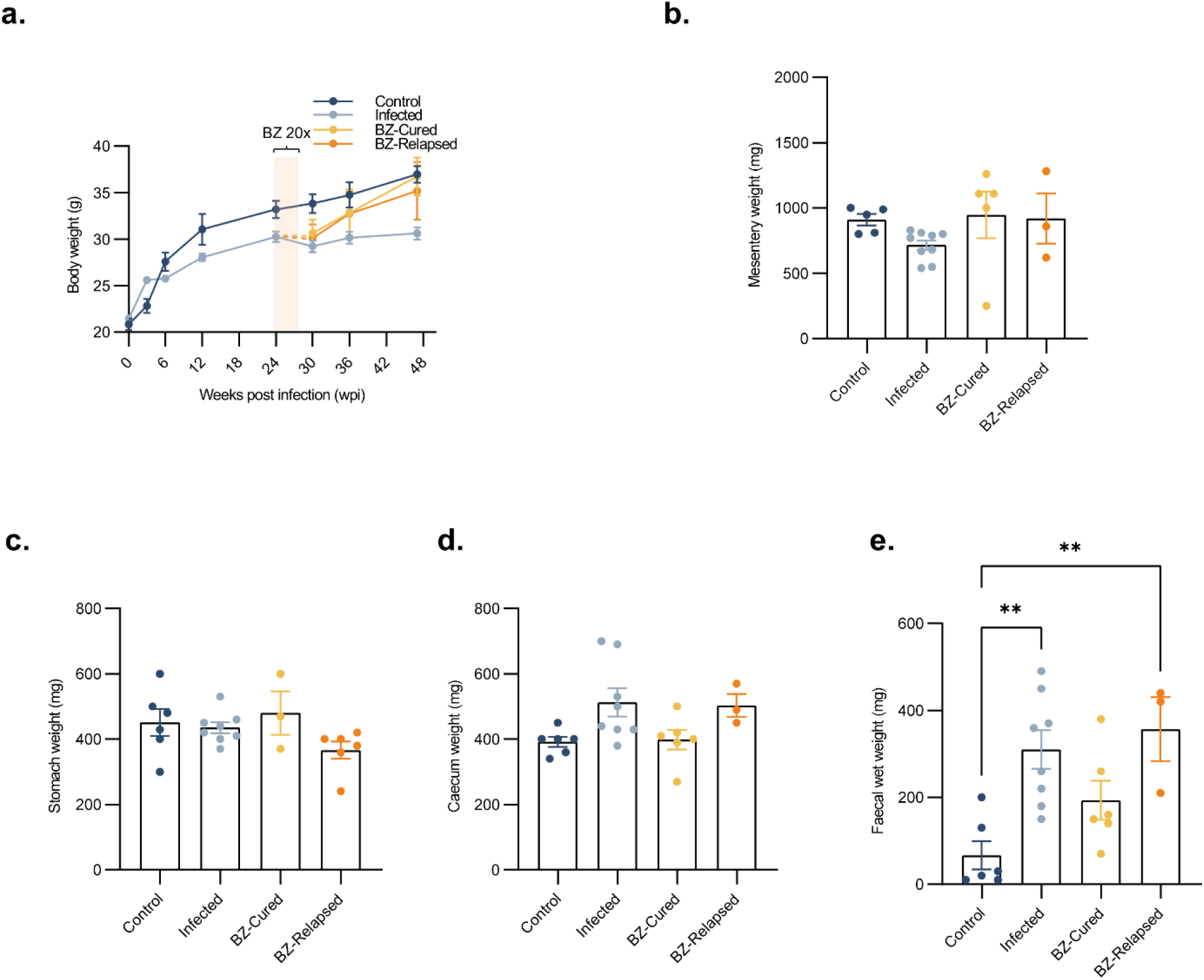
Additional GI assays after benznidazole treatment initiated at 24 weeks post-infection. **a**, Line plots show body weight of control (*n* = 5, except *n* =10 at 0, 3 and 24 weeks post-infection (wpi)), infected (*n* = 9, except *n =* 25 at 0 and 3 wpi, *n* = 20 at 6 wpi, *n* = 23 at 12 wpi, and *n* = 22 at 24 wpi), benznidazole treated cured (BZ-Cured; *n* = 7) and benznidazole treated relapsed (BZ-relapsed; *n* = 3) mice. Overlaid cream bar shows the BZ treatment window (24 - 27 wpi, 20 daily doses). Bar plots show **b**, mesentery, **c**, stomach **d**, caecum and **e**, faecal wet weight of control (*n* = 5), infected (*n* = 8), BZ-Cured (*n* = 3 - 6) and BZ-Relapsed (*n* = 3) mice at 36 wpi. Statistical significance was tested using one-way ANOVA followed by Tukey’s HSD test (\*\**P* < 0.01).

**Extended Data Figure 6:**
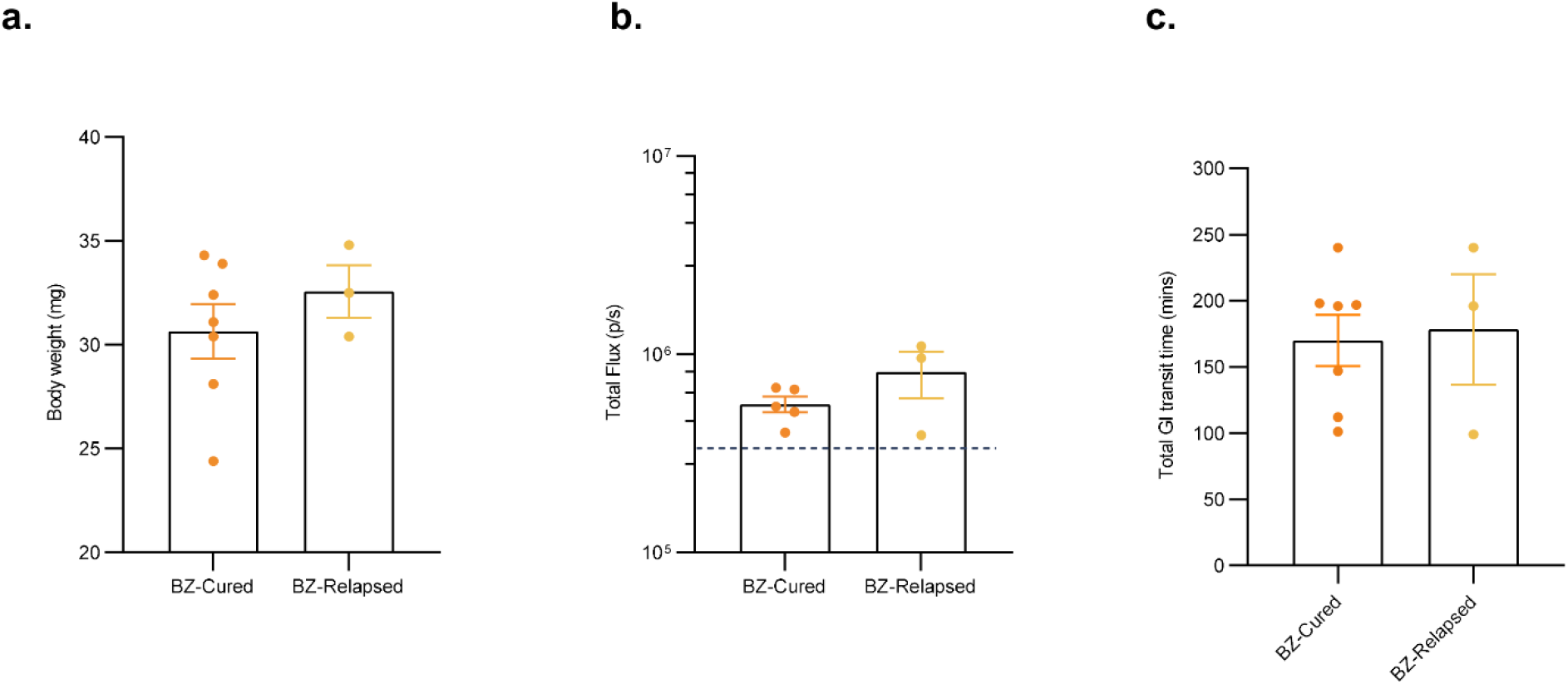
Pre-treatment characteristics of relapses compared with cures for benznidazole treatment initiated at 24 weeks post-infection. Bar plots show characteristics of benznidazole treated and cured (BZ-Cured, *n* = 7) and benznidazole treated and relapsed (BZ-Relapsed, *n* = 3) mice retrospectively segregated at the point of treatment initiation (24 weeks postinfection (wpi)): **a**, body weight; **b**, bioluminescence intensity (total flux) and **c**, total GI transit time.

**Supplementary Table 1:**
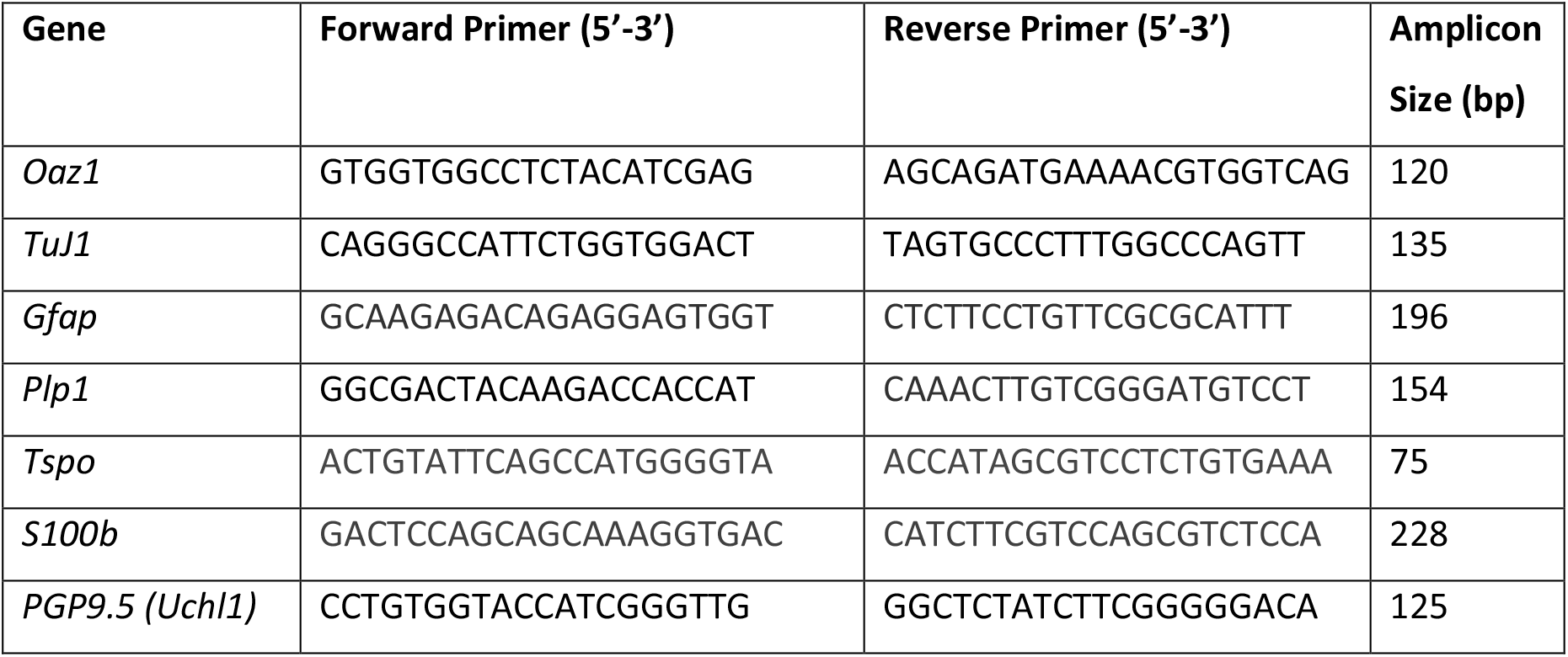
qPCR experiment primer design.

